# Membrane binding by CHMP7 coordinates ESCRT-III dependent nuclear envelope reformation

**DOI:** 10.1101/049221

**Authors:** Yolanda Olmos, Anna Perdrix, Jeremy G Carlton

## Abstract

Amongst other cellular functions, the Endosomal Sorting Complex Required for Transport-III (ESCRT-III) machinery controls nuclear envelope (NE) reformation during mitotic exit by sealing holes in the reforming NE. ESCRT-III also acts to repair this organelle upon migration-induced rupture. The ESCRT-III component CHMP7 is responsible for recruitment of ESCRT-III to the NE. Here, we show that the N-terminus of CHMP7, comprising tandem Winged Helix (WH)-domains, is a membrane-binding module. This activity allows CHMP7 to bind to the Endoplasmic Reticulum (ER), an organelle continuous with the NE, and provides a platform to direct NE-recruitment of ESCRT-III during mitotic exit. Point mutations that disrupt membrane-binding prevent CHMP7 localising to the ER and its subsequent enrichment at the reforming NE. These mutations prevent both assembly of downstream ESCRT-III components at the reforming NE and proper establishment of post-mitotic nucleo-cytoplasmic compartmentalisation. These data identify a novel membrane-binding activity within an ESCRT-III subunit that is essential for post-mitotic nuclear regeneration.

**One Sentence Summary:** CHMP7’s atypical N-terminus is a membrane-binding module that allows assembly and function of ESCRT-III at the nuclear envelope during mitotic exit.

## Introduction

ESCRT-III regulates an array of topologically equivalent membrane remodelling events including multivesicular body (MVB) biogenesis (Babst *et al*, 2002), release of enveloped retroviruses including HIV-1 (Martin-Serrano & Neil, 2011), abscission during cytokinesis (Carlton & Martin-Serrano, 2007; Morita *et al*, 2007), repair of the plasma membrane (Jimenez *et al*, 2014; Scheffer *et al*, 2014), post-mitotic reformation of the NE (Olmos *et al*, 2015; Vietri *et al*, 2015) and repair of the NE under pathological conditions (Raab *et al*, 2016; Denais *et al*, 2016). In these processes, ESCRT-III is thought to function on the cytoplasmic face of a membranous stalk, acting to resolve this stalk, and bringing about separation of previously connected membranes. In yeast, ESCRT-III also surveilles and extracts damaged nucleoporins from the NE (Webster *et al*, 2014). ESCRT-III is a filament-forming complex comprising polymers of CHMPs (*Ch*arged *M*ultivesicular Body *P*roteins/*Ch*ro*m*atin Remodelling *P*roteins). CHMPs are soluble cytoplasmic proteins, but can transition to a filament-forming state upon co-polymerisation. Adaptors and upstream ESCRT components (ESCRT-I, ESCRT-II and ESCRT-associated proteins such as ALIX or HDPTP) recruit ESCRT-III to sites of activity; during cytokinesis, CEP55 recruits ESCRT-III via TSG101 and ALIX; during plasma membrane repair, ALG2 is necessary for ESCRT-III recruitment (Scheffer *et al*, 2014); during HIV-1 release, viral Gag proteins recruit ESCRT-III via TSG101 and ALIX; during MVB biogenesis, an HRS/Vps27-containing complex recruits ESCRT-III to endosomes via ESCRT-I and ESCRT-II. Once recruited, ESCRT-III assembles in a defined order, with CHMP4 proteins recruiting CHMP2 and CHMP3 proteins (Carlton, 2010; McCullough *et al*, 2013; Hurley, 2015; Teis *et al*, 2008; Henne *et al*, 2012). In the context of NE reformation, the ESCRT-III complex assembles in a classical order and whilst factors such as UFD1 stimulate incorporation of late-acting ESCRT-III components (Olmos *et al*, 2015), the machinery itself is recruited by the poorly studied ESCRT-III subunit, CHMP7 (Vietri *et al*, 2015; Denais *et al*, 2016). Here, we set out to discover activities in CHMP7 that contribute to ESCRT-III function on the reforming NE.

## Results and Discussion

CHMP7 is unique amongst ESCRT-III subunits in that it contains an extended N-terminus (NT) (Figure S1A) that we hypothesised may be important for its role in NE regeneration. To minimise artefacts associated with fixation and to preserve signal intensity when imaging, we imaged living cells stably-expressing low levels of GFP-CHMP7 expressed from weak retroviral promoters. We were surprised to find that whilst CHMP7 was recruited as expected to the NE during mitotic exit, in addition to a cytoplasmic pool, it decorated ER membranes in interphase and mitotic cells (Figure 1A-Figure 1C, Figure S1B-S1D, Movie 1-4). Whilst antibodies against CHMP7 were unsuitable for immunofluorescence, we could detect endogenous CHMP7 in ER-fractions from homogenised cells (Figure 1D, Figure 1E and Figure S1E) and saw no localisation of CHMP7 to the midbody (Figure S1F). We developed siRNA targeting CHMP7 and visualised ER localisation and NE enrichment in CHMP7-depleted cell lines stably expressing near endogenous levels of siRNA-resistant GFP-CHMP7 (Figure S2, Movie 5). Long thought to be absent from yeast, *S. cerevisiae* Chm7 was recently shown to localise to the ER (Bauer *et al*, 2015), suggesting that this localisation is evolutionarily conserved. During NE reformation, all other ESCRT-III subunits are recruited from the cytoplasm (Olmos *et al*, 2015; Vietri *et al*, 2015); given that the NE is formed from the ER (Lu *et al*, 2009; Anderson & Hetzer, 2007), a pre-existing ER localisation for CHMP7 suggests a platform from which this recruitment could occur. Analysis of stable cell lines stably expressing GFP-CHMP7^NT^ revealed that CHMP7’s N-terminus directed localisation to the ER (Figure 1A, Movie 6, Movie 7), but this protein exhibited little stabilisation at the reforming NE (Figure S3A). ER-localisation was confirmed in cells stably expressing mCh-CHMP7^NT^ (Fig S3B, S3C). In contrast, the C-terminus of CHMP7 (CHMP7 δNT) was cytosolic and displayed neither ER-localisation nor stabilisation at the reforming nuclear envelope (Figure 1F, Movie 8), despite containing the CHMP4B/ESCRT-III interaction domain(Horii *et al*, 2006). CHMP7 is responsible for recruiting downstream ESCRT-III components to the reforming NE through CHMP4B (Figure S3D-S3F) (Vietri *et al*, 2015). Fusion of siRNA-resistant CHMP7^NT^ to CHMP4B directed cytoplasmic CHMP4B to the mitotic ER and restored its enrichment at sites of annular fusion at the forming NE in the absence of endogenous CHMP7 (Figure S3B-S3D). These data indicate that CHMP7’s NT is an ER-localisation domain that is essential for downstream ESCRT-III assembly at the reforming NE.

**Figure 1:**
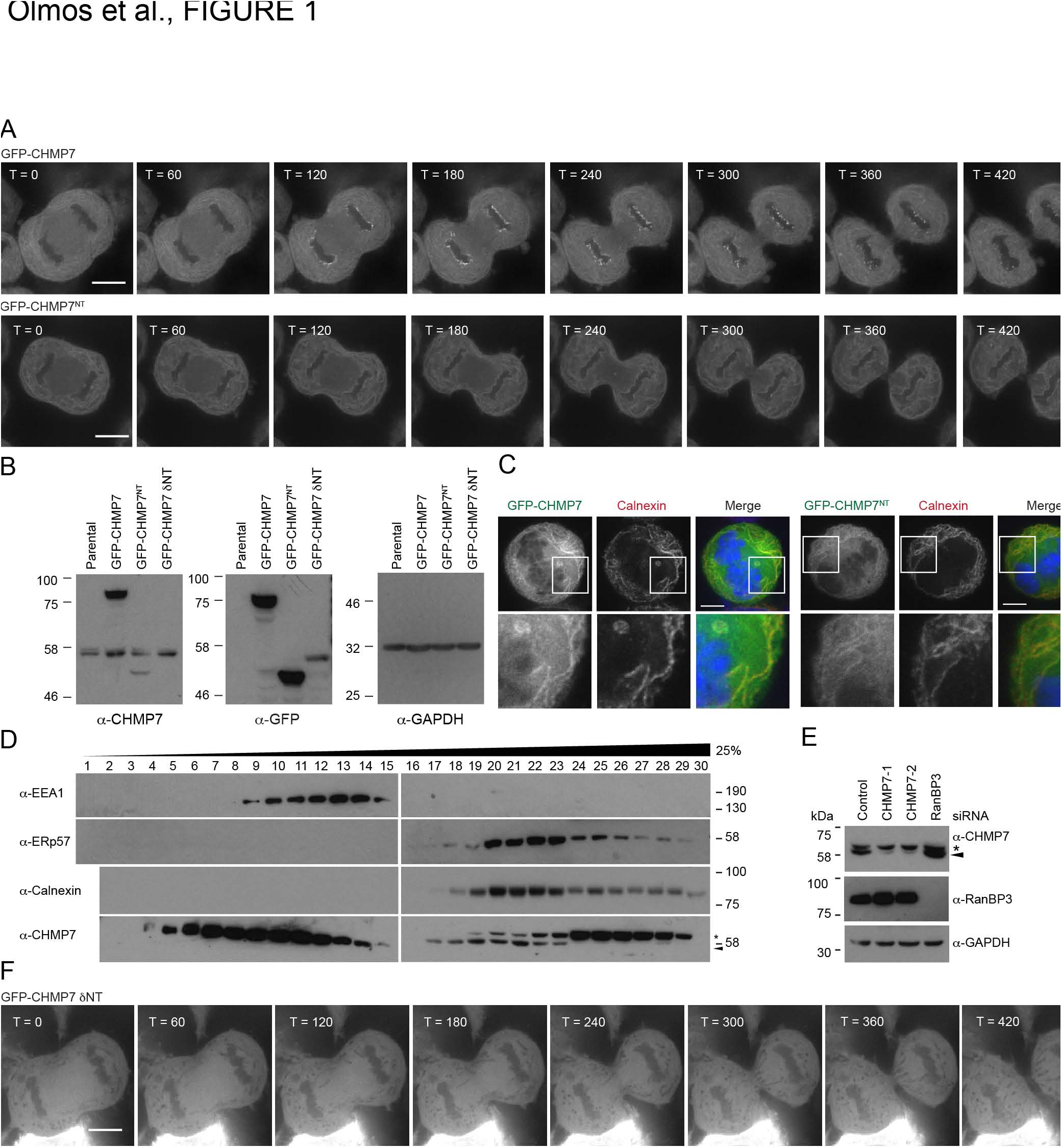
Mapping activities in CHMP7 that govern ER and NE localisation. A-C. HeLa cells stably expressing GFP-CHMP7 or GFP-CHMP7^NT^ were imaged live (A), lysed, resolved and examined by western blotting with anti-GFP, anti-CHMP7 or anti-GAPDH antisera (B) or fixed and stained with anti-calnexin antisera and 4′,6-diamidino-2-phenylindole (DAPI) (C). Scale bar is 10 μm. Images in A are representative of all cells imaged and 22/22 (GFP-CHMP7) and 21/21 (GFP-CHMP7^NT^) captured movies, co-localisation of GFP-CHMP7 and GFP-CHMP7^NT^ with Calnexin was observed in 7/7 and 13/13 scored cells respectively. Time given in seconds post cortical ingression. D. Post-nuclear supernatants from Cos7 cells were fractionated through a continuous iodixanol gradient and analysed by SDS-PAGE and western blotting with the indicated antisera. * is a non-specific band, endogenous CHMP7 indicated by arrowhead. E. Resolved cell lysates from HeLa cells transfected with the indicated siRNAs were examined by western blotting with anti-CHMP7, anti-GAPDH or anti-RanBP3 antisera, * is a non-specific band, endogenous CHMP7 indicated by arrowhead. F. HeLa cells stably expressing GFP-CHMP7 δNT were imaged live. Presented images are representative of all cells imaged and 5/5 captured movies. Scale bar is 10 μm.

Analysis of the secondary structure of CHMP7^NT^ has revealed the presence of tandem winged helix (WH) domains (Sundquist & Ullman, 2015; Bauer *et al*, 2015) resembling those found in ESCRT-II subunits (Figure S4A). In the context of endosomal-sorting, membrane anchored ESCRT-II serves to recruit ESCRT-III to endosomes through interaction of the 2^nd^ WH domain of VPS25 with the ESCRT-III component VPS20/CHMP6 (Teis *et al*, 2010; Im *et al*, 2009). Bauer et al. (Bauer *et al*, 2015) have suggested that CHMP7 represents a fusion of ESCRT-II and ESCRT-III subunits, and given the role for CHMP7 in initiating ESCRT-III assembly at the NE, we wondered if this region of CHMP7 acted as a membrane-adaptor at this organelle. HH-Pred alignments of CHMP7 matched its NT to VPS25 (Bauer *et al*, 2015) and by aligning predicted secondary structural elements in CHMP7 to those present in the crystal structure of VPS25, we noted an evolutionarily conserved extension of the loop between the β2-β3 hairpin in the 1^st^ WH-domain of CHMP7^NT^ (Figure S4A).

As deletions through CHMP7^NT^ destabilised the protein (Fig S3B), we performed scanning mutagenesis through CHMP7^NT^ to identify regions necessary for ER-localisation (Figure S4B, S4C). We discovered 12 mutagenic tetrads that prevented ER localisation, five of which were found on the extended loop in the β2-β3 hairpin in the 1^st^ WH-domain of CHMP7^NT^. To understand where the remainder lay, we created a homology model of CHMP7^NT^ (lacking the extended loop between WH1 β2-β3) based upon the crystal structure of VPS25 (Figure S4D). This model revealed that the remainder of mutations mapped to regions that were either in, or engaged with, residues on the β2-β3 hairpin in the 1^st^ WH-domain of CHMP7^NT^ (Figure S4D). Replacement of the loop between the β2-β3 hairpin with a Gly-Ser-Gly-Ser linker (δ107-148) prevented ER localisation (Figure 2A). As alanine changes in blocks of 4 may have detrimental effects on the secondary structure of CHMP7^NT^, we mutated individual residues within this loop to discover if more conservative mutations could disrupt ER localisation. This revealed that mutation of 6 evolutionarily conserved hydrophobic residues (W118, W121, F126, L127, L128 and L131, or deletion of this hydrophobic stretch (δ118-128, Figure 2A and 2B) prevented ER localisation of CHMP7^NT^. We introduced these mutations in the context of full-length CHMP7 and found that the point mutations and deletions that prevented ER-localisation of GFP-CHMP7^NT^ also prevented localisation of full-length GFP-CHMP7 to this membrane and further prevented subsequent enrichment of this protein at the reforming NE (Figure 2C, Movies 9-14). These data indicate that CHMP7 cannot be recruited to the reforming NE from the cytoplasm.

**Figure 2:**
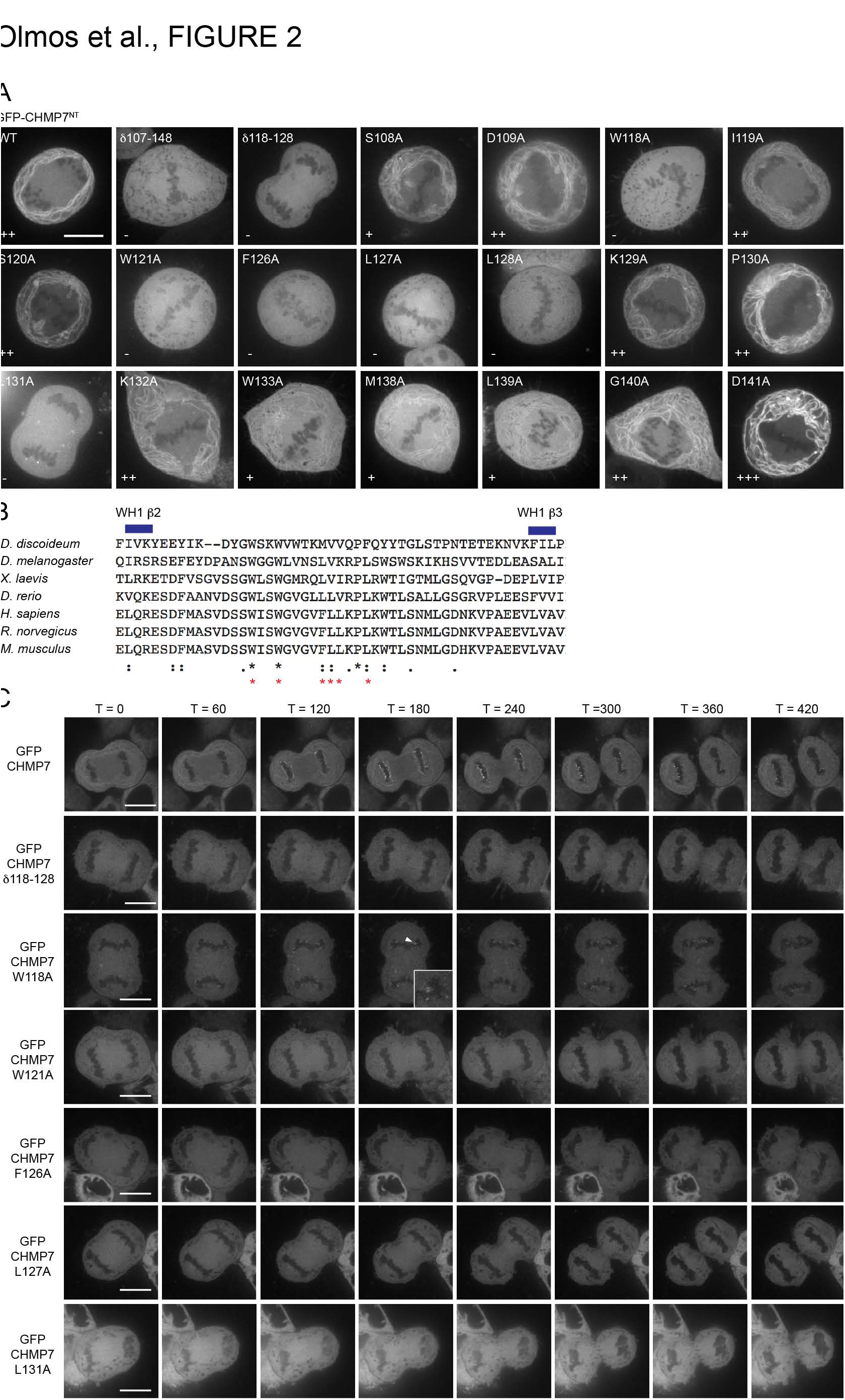
Mapping activities in CHMP7^NT^ that govern ER localisation. A. HeLa cells were transfected with the indicated GFP-CHMP7^NT^ plasmids and imaged live; individual residues within the β2-β3 insertion that disrupted ER-localisation when previously mutated in blocks of 4 (Fig S4B-D) were mutated to alanine. Scale bar is 10 μm. Images representative of 3/3 acquired images per mutation. B. Sequence alignment of insertion between β2 and β3 in the CHMP7^NT^ WH1 domain from the indicated organisms. C. HeLa cells expressing indicated GFP-CHMP7 constructs were imaged live. Scale bar is 10 μm, time in seconds post cortical ingression presented. Images are representative of 3/3 acquired movies and the cytoplasmic nature was observed in 50/50 scored cells per mutation. Limited enrichment on the telophase NE (boxed) was observed for GFP-CHMP7 W118A, suggesting that some degree of ER-localisation persists in this case.

Given the requirement for conserved hydrophobic residues in establishing ER-localisation, we wondered if they acted as a membrane-binding region to anchor this protein in the ER. In support of this, we found that GST-CHMP7^NT^, but not GST, could be avidly captured upon liposomes (Figure 3A-D). HIS-CHMP7^NT^ could also be captured on liposomes (Figure S5A), excluding the possibility that the GST-tag was influencing binding. The fusogenic lipid diacylglycerol has been implicated in NE reformation (Domart *et al*, 2012), however, we found membrane interaction of CHMP7^NT^ was insensitive to the presence of diacylglycerol (Figure 3C, 3D). We also found membrane interaction to be insensitive to the degree of membrane-curvature (Figure S5B, S5C). Mutation of residues that disrupted ER-localisation prevented GST-CHMP7^NT^ from binding liposomes, with deletion of the hydrophobic cluster (δ118-128), or individual mutation of L127A or L131A having the strongest effect (Figure 3E and 3F). Importantly, these mutations did not destabilise GST-CHMP7^NT^ (Figure S5D). These data indicate that an evolutionarily conserved cluster of hydrophobic amino acids in an extended loop on the 1^st^ WH-domain of CHMP7^NT^ act as a membrane anchor that directs ER-localisation and provides a platform for ESCRT-III recruitment at the reforming NE.

**Figure 3:**
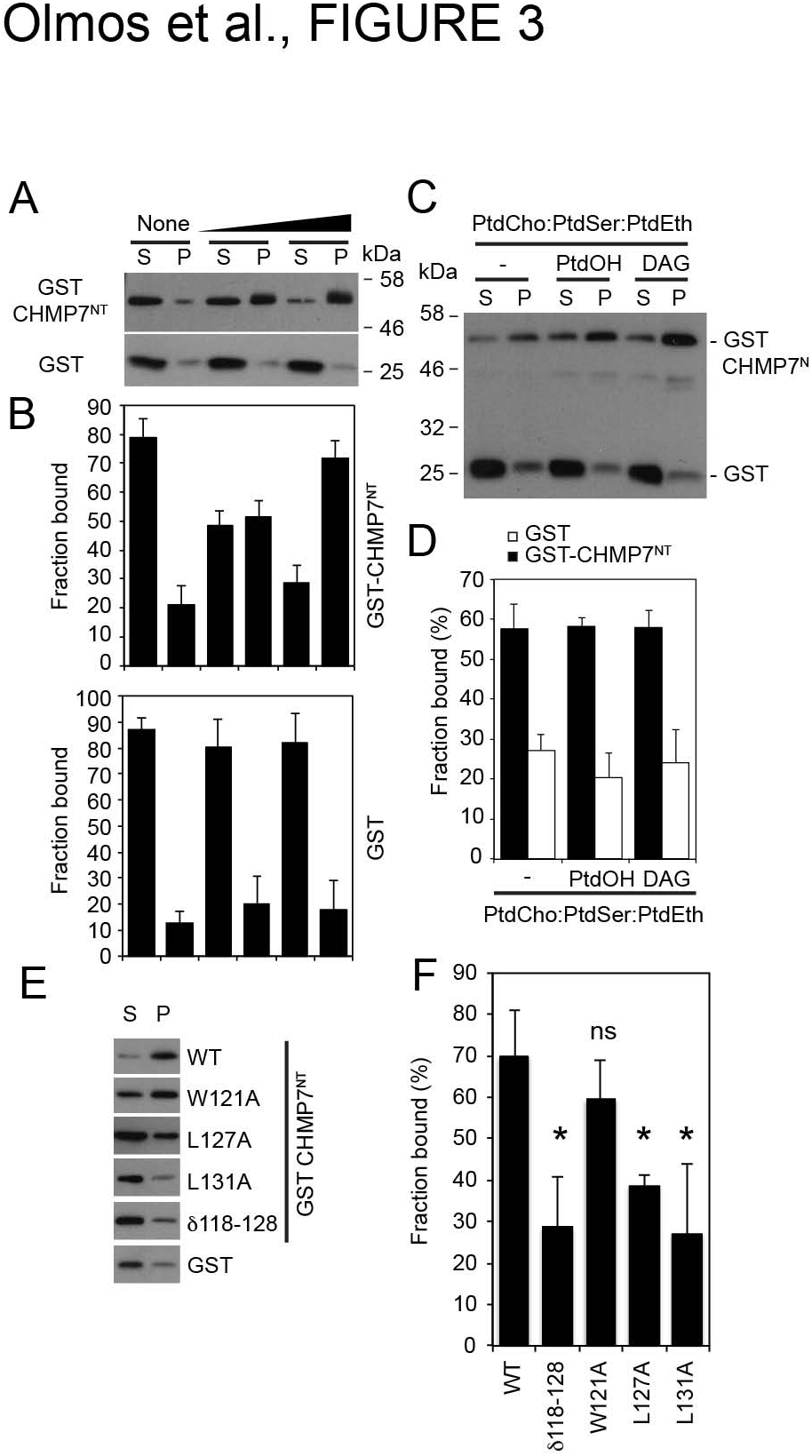
CHMP7^NT^ binds lipid membranes. A-D. GST or GST-CHMP7^NT^ was incubated for 5 minutes (A) or 15 minutes (B, C) with Folch (A, B) or synthetic (C, D) liposomes. Liposomes were collected by ultracentrifugation. Pelleted (P) and soluble (S) fractions were resolved, analysed by western blotting with anti-GST antisera and quantified by densitometry; captured fraction presented in (B and D) as fraction bound. E. GST-CHMP7^NT^ bearing the indicated mutations was incubated with Folch liposomes and liposome binding was assessed as previously described. WT, N = 7; δ118-128, N = 7; W121A, N = 7, ns; L127A, N = 4; L131A, N = 4. Statistical significance calculated using 1-way ANOVA with Dunnett’s multiple comparison test, * = P <0.0001.

Consistent with a role for ESCRT-III in ensuring post-mitotic nucleocytoplasmic compartmentalisation (Olmos *et al*, 2015; Vietri *et al*, 2015) and a role for CHMP7 in recruiting CHMP4 proteins to the reforming NE (Figure S3B – S3D) (Vietri *et al*, 2015), we found that CHMP7 depletion prevented enrichment of endogenous CHMP2A at the reforming NE (Figure 4A and 4B). Failure to recruit CHMP2A to this organelle has been shown to leave holes in the reforming NE (Olmos *et al*, 2015) and, consistent with this, we found that CHMP7 depletion leads to a poorly sealed post-mitotic nuclear envelope (Figure 4C and 4D). Assembly of endogenous CHMP2A at the telophase NE could be rescued by stable expression of siRNA-resistant HA-CHMP7 (HA-CHMP7^R^) or HA-CHMP7^NT-R^/CHMP4B, expressed at near endogenous levels, but could not be supported in CHMP7-depleted cells stably expressing HA-CHMP7^R^ δNT, HA-CHMP7^R^ δ118-128, or HA-CHMP7^R^ L127A (Figure 4E and 4F). Notably, midbody accumulation of endogenous CHMP2A was unaffected in CHMP7-depleted cells expressing HA-CHMP7^R^ δNT (Figure 4G), indicating that the membrane-binding ability of CHMP7 is specifically required for ESCRT-III assembly at the NE. Further, whilst the nucleocytoplasmic compartmentalisation defect induced by CHMP7-depletion could be rescued by stable expression of HA-CHMP7^R^ or HA-CHMP7^NT-R^/CHMP4B, stable expression of HA-CHMP7^R^ δNT, HA-CHMP7^R^ δ118-128, or HA-CHMP7^R^ L127A failed to rescue this compartmentalisation defect (Figure 4H and 4I).

**Figure 4:**
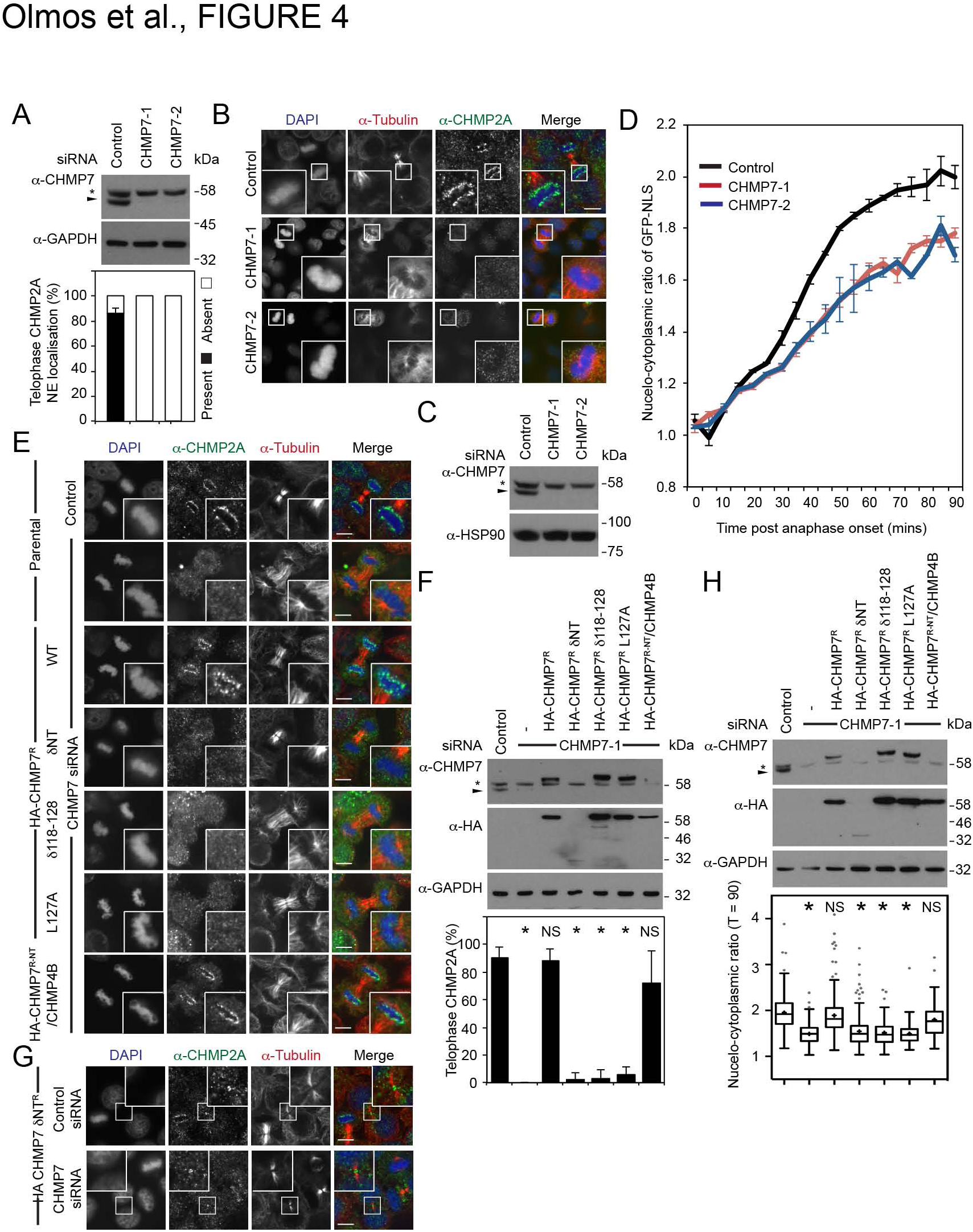
Membrane binding by CHMP7 is essential for ESCRT-III-dependent NE reformation during mitotic exit. A, B. Resolved lysates of HeLa cells transfected with the indicated siRNA were examined by western blotting with anti-CHMP7 or anti-GAPDH antisera (A). Cells were fixed, stained with anti-tubulin, anti-CHMP2A and DAPI and examined by immunofluorescence, scale bar is 10 μm (B). Assembly of ESCRT-III at the telophase NE was quantified (A; Bars represent mean ± S.D., n = 40, N = 2, P = 0.0008, as calculated by two-tailed Student’s T-test). C, D. Resolved lysates of HeLa cells stably expressing GFP-NLS and H2B-mCh and transfected with the indicated siRNA, were analysed by western blotting with anti-CHMP7 and anti-GAPDH antisera (C). Cells were imaged live and the degree of nucleocytoplasmic compartmentalisation was calculated at the indicated timepoints (D). E, F. HeLa cells stably expressing the indicated HA-tagged, siRNA-resistant CHMP7 proteins were transfected with control or CHMP7-targeting siRNA, fixed, stained with anti-tubulin, anti-CHMP2A and DAPI and examined by immunofluorescence, scale bar is 10 μm (E) and quantified (F; Control, N = 5, n = 80; CHMP7 siRNA, N = 4, n = 34, P < 0.001; CHMP7 siRNA and HA-CHMP7^R^, N = 4, n = 52, NS (P = 0.995); CHMP7 siRNA and HA-CHMP7^R^ δNT, N = 4, n = 40, P < 0.001; CHMP7 siRNA and HA-CHMP7^R^ δ118-128, N = 3, n = 30, P < 0.001; CHMP7 siRNA and HA-CHMP7^R^ L127A, N = 3, n = 32, P < 0.001; CHMP7 siRNA and HA-CHMP7^R-NT^/CHMP4B, N = 3, n = 31, NS (P = 0.055)). Graphs present mean ± S.D. from the indicated number (N) of independent experiments. Statistical significance calculated using 1-way ANOVA with Dunnett’s multiple comparison test, * = significant. Resolved lysates of cells from E were analysed by western blotting with anti-CHMP7, anti-HA and anti-GAPDH antisera, endogenous CHMP7 marked by arrowhead, * is a non-specific band (F). G. HeLa cells stably expressing HA-CHMP7^R^ δNT were transfected with control or CHMP7-targetting siRNA, fixed, stained with anti-tubulin, anti-CHMP2A and DAPI and examined by immunofluorescence, scale bar is 10 μm. Midbody localisation observed in 30/30 cases (Ctrl) and 29/30 cases (CHMP7 siRNA) N = 3. H. Resolved lysates of HeLa cells stably expressing GFP-NLS, H2B-mCh and the indicated HA-tagged, siRNA-resistant CHMP7 proteins were transfected with control or CHMP7-targeting siRNA and examined by western blotting with anti-CHMP7, anti-HA or anti-GAPDH antisera, endogenous CHMP7 marked by arrowhead, * is a non-specific band. Cells were imaged live and the degree of nucleocytoplasmic compartmentalisation was calculated 90 minutes post anaphase-onset (Control, 1.93 ± 0.04, N = 9, n = 236; CHMP7 siRNA, 1.51 ± 0.03, N = 9, n = 252, P < 0.0001; CHMP7 siRNA and HA-CHMP7^R^, 1.88 ± 0.05, N = 8, n = 257, NS (P = 0.8909); CHMP7 siRNA and HA-CHMP7^R^ δNT, 1.54 ± 0.08, N = 4, n = 207, P = < 0.0001; CHMP7 siRNA and HA-CHMP7^R^ δ118-128, 1.53 ± 0.1, N = 3, n = 101, P = 0.0003; CHMP7 siRNA and HA-CHMP7^R^ L127A, 1.50 ± 0.08, N = 3, n = 101, P = 0.0001; CHMP7 siRNA and HA-CHMP7^R-NT^/CHMP4B, 1.73 ± 0.1, N = 3, n = 76, NS (P = 0.1556). Values quotes as mean ± S.E.M. from the indicated number (N) of independent experiments containing the indicated number (n) of measured cells. Statistical significance calculated using 1-way ANOVA with Dunnett’s multiple comparison test from experimental means (N), * = significant, Tukey whiskers displayed, means marked by xs+).

We describe a novel localisation of CHMP7 to the ER and its subsequent function in regenerating a sealed nuclear envelope during mitotic exit. Given the continuity of the ER with the NE (Burke & Ellenberg, 2002), this localisation provides a mechanistic understanding for how cytosolic ESCRT-III subunits can be recruited to, and assemble upon, this organelle. Whilst the N-terminus of CHMP7 could not be stabilised at sites of annular fusion, it is likely that subsequent engagement of ESCRT-III and partners by the C-terminus of CHMP7 provides this stabilising cue (Vietri *et al*, 2015; Olmos *et al*, 2015; Horii *et al*, 2006). CHMP7^NT^ comprises tandem WH-domains that resemble the ESCRT-II subunit VPS25, and consistent with a role for ESCRT-II in recruiting ESCRT-III to cellular membranes, this region directs ESCRT-III assembly at the NE. WH-1 of CHMP7^NT^ contains a membrane anchor that localises this protein in the ER; *C. elegans* ESCRT-II has been suggested to localise to the sarcoplasmic reticulum, suggesting that the tandem WH-fold may play a broader role in ER targeting (Lefebvre *et al*, 2016). Membrane binding by the CHMP7^NT^ domain is necessary for ER localisation, subsequent enrichment of CHMP7 at the reforming NE and, given CHMP7’s ability to bind CHMP4 proteins (Horii *et al*, 2006), is essential for assembly of downstream ESCRT-III components at sites of annular fusion. Importantly, we demonstrate that these ESCRT-III subunits cannot be recruited to sites of annular fusion from the cytosol in the absence of a membrane anchored CHMP7 to facilitate their incorporation at this organelle. CHMP7 is thus an ER-specific membrane-adaptor for ESCRT-III that provides an activity essential for post-mitotic organelle biogenesis and maybe necessary for repair of the nuclear envelope under physiological (Vargas *et al*, 2012) and pathological (Raab *et al*, 2016; Denais *et al*, 2016) conditions.

## Author Contributions

Conception and design: JGC. Acquisition, analysis, interpretation of data: JGC, AP and YO. Drafting and revising manuscript: JCG and YO.

## Acknowledgements

JCG is a Wellcome Trust Research Career Development Fellow (093603/Z/10/Z). We acknowledge the Nikon Imaging Centre at KCL for access to core equipment. We thank Camille Wouters and Nisreen Chahid who assisted with cloning as part of high-school Nuffield Research Placements. We thank Dr Pierfrancesco Marra (KCL) for guidance on density gradient centrifugation.

## Conflict of Interest

The authors declare that they have no conflicts of interest

## Supplemental Figure Legends

**Figure S1:**
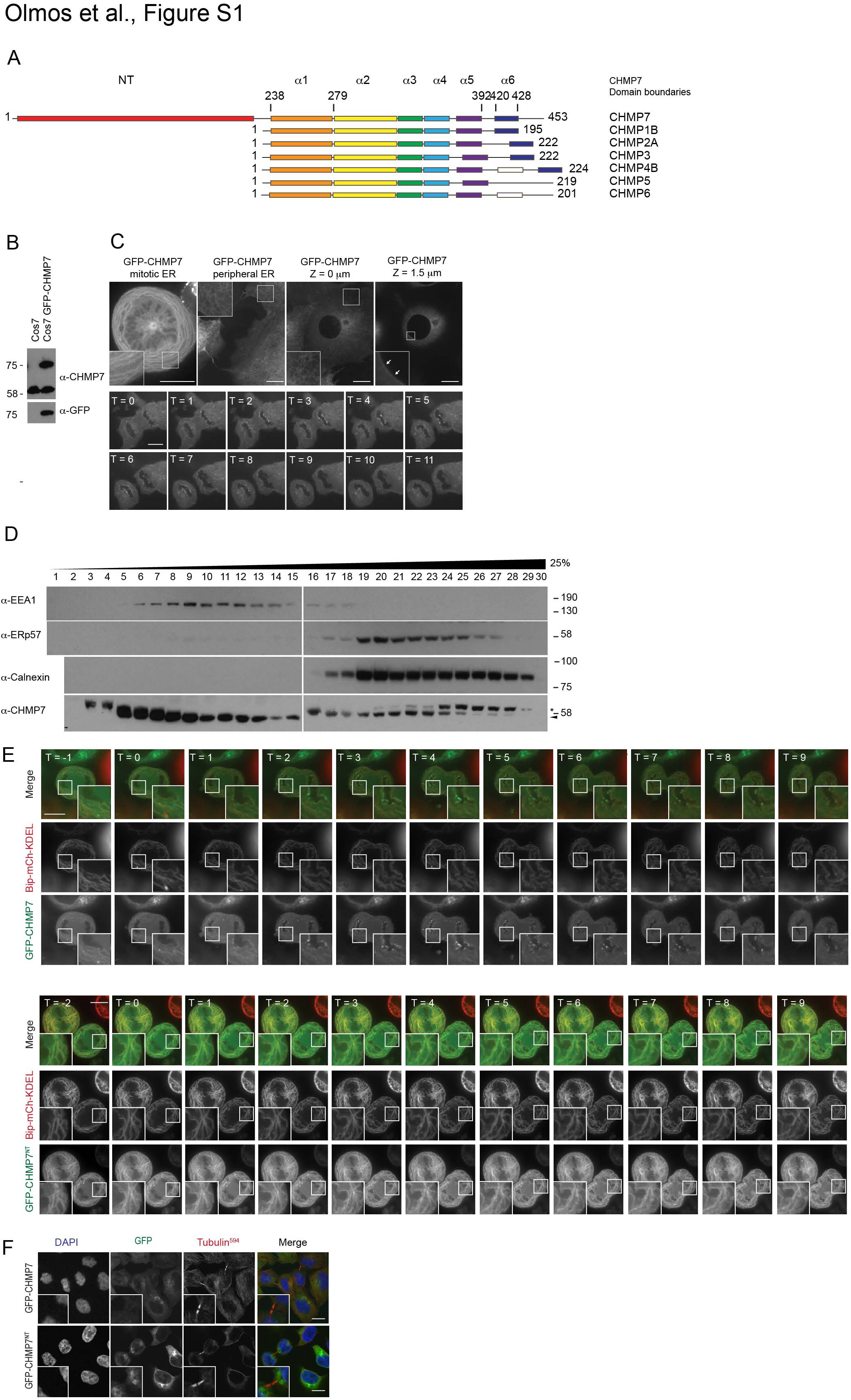
CHMP7 localises to the ER. A. Domain analysis of CHMP7, diagram adapted from Bajorek et al., 2009(Bajorek *et al*, 2009). B, C. Resolved cell lysates of Cos7 cells and Cos7 stably expressing GFP-CHMP7 at endogenous levels were analysed by western blotting with anti-CHMP7 and anti-GFP antisera (B) or imaged live (C). Scale bar is 10 μm, images representative of all cells imaged and 72/72 captured interphase cells, 11/11 captured mitotic cells and 5/5 captured movies of enrichment of GFP-CHMP7 at the telophase NE. Note, GFP-CHMP7 was present on the interphase NE (arrows), but unlike during mitotic NE reformation, was not enriched upon this membrane compared to the ER. ER-localisation of GFP-CHMP7^NT^ was observed in all visualised and 19/19 captured interphase HeLa cells (data not shown). D. Post-nuclear supernatants from HeLa cells were fractionated through a continuous iodixanol gradient and analysed by SDS-PAGE and western blotting with the indicated antisera. * is a non-specific band, endogenous CHMP7 indicated by arrowhead. E. HeLa cells stably expressing GFP-CHMP7 or GFP-CHMP7^NT^ were transfected with pLHCX BIP-mCh-KDEL and imaged live. Images representative of all cells imaged and 3/3 captured movies per condition. Scale bar is 10 μm. F. HeLa cells stably expressing GP-CHMP7 or GP-CHMP7^NT^ were fixed and stained with anti-tubulin and DAP. Scale bar is 10 μm.

**Figure S2:**
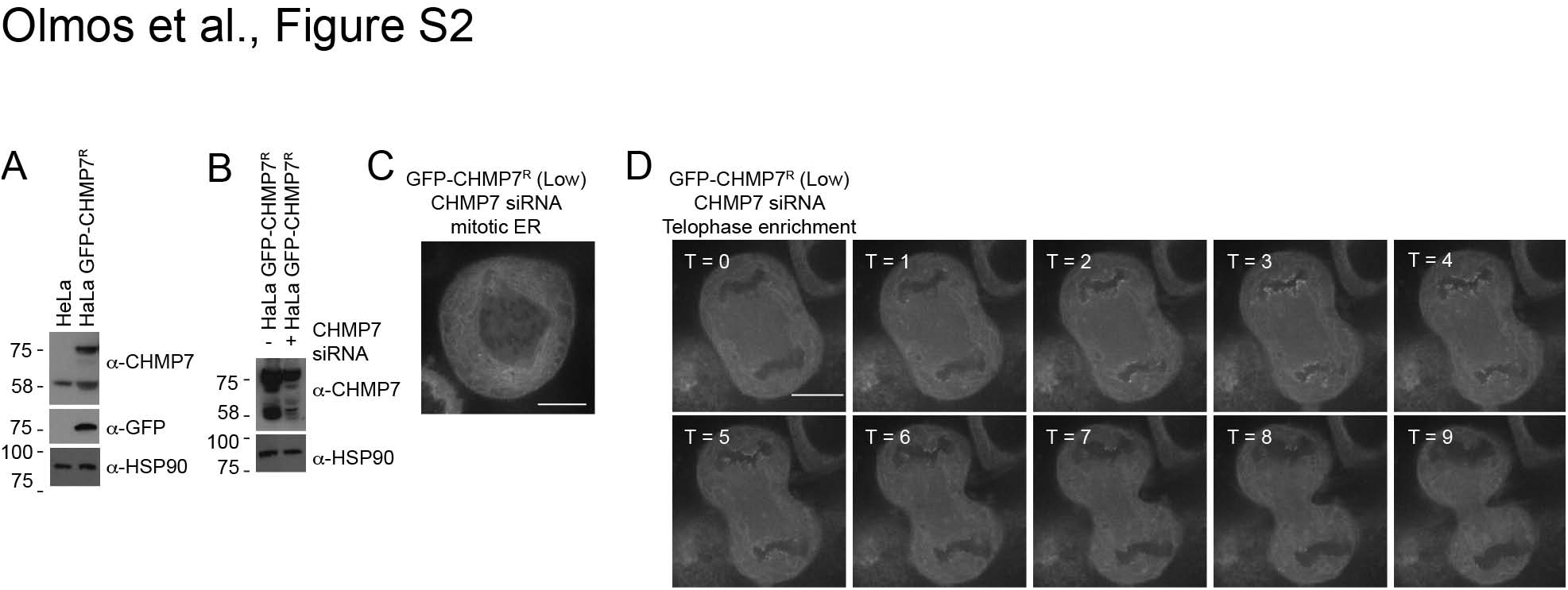
GFP-CHMP7 localisation at endogenous expression levels. A, B. Resolved lysates of HeLa cells or HeLa cells stably expressing GP-CHMP7^R^ from a weaker retroviral vector (pNG72) were transfected with siRNA as indicated (B) and analysed by western blotting with anti-CHMP7, anti-GFP or anti-HSP90 antisera (A, B). Alternatively, siRNA-transfected cells were imaged live (C, D). Whilst the HeLa cells presented in Figure 1A express GFP-CHMP7 in excess of endogenous CHMP7, ER-localisation and NE-enrichment of GFP-CHMP7 was observed in these CHMP7-depleted HeLa cells stably expressing siRNA-resistant GFP-CHMP7 at near endogenous levels (1.6 ± 0.2 fold, n = 3 ± S.D.; reticular localisation to mitotic ER observed in all imaged cells and 38/38 acquired cells; enrichment at the reforming NE observed in 7/7 acquired movies). Imaging in this case was technically challenging due to the very low level of GFP-signal at this expression level, to facilitate image acquisition and to ensure inferences of localisation defects were not compounded by poor-signal strength, we elected to use cells expressing higher GFP-CHMP7 for subsequent mutagenic analysis in Figure 2C.

**Figure S3:**
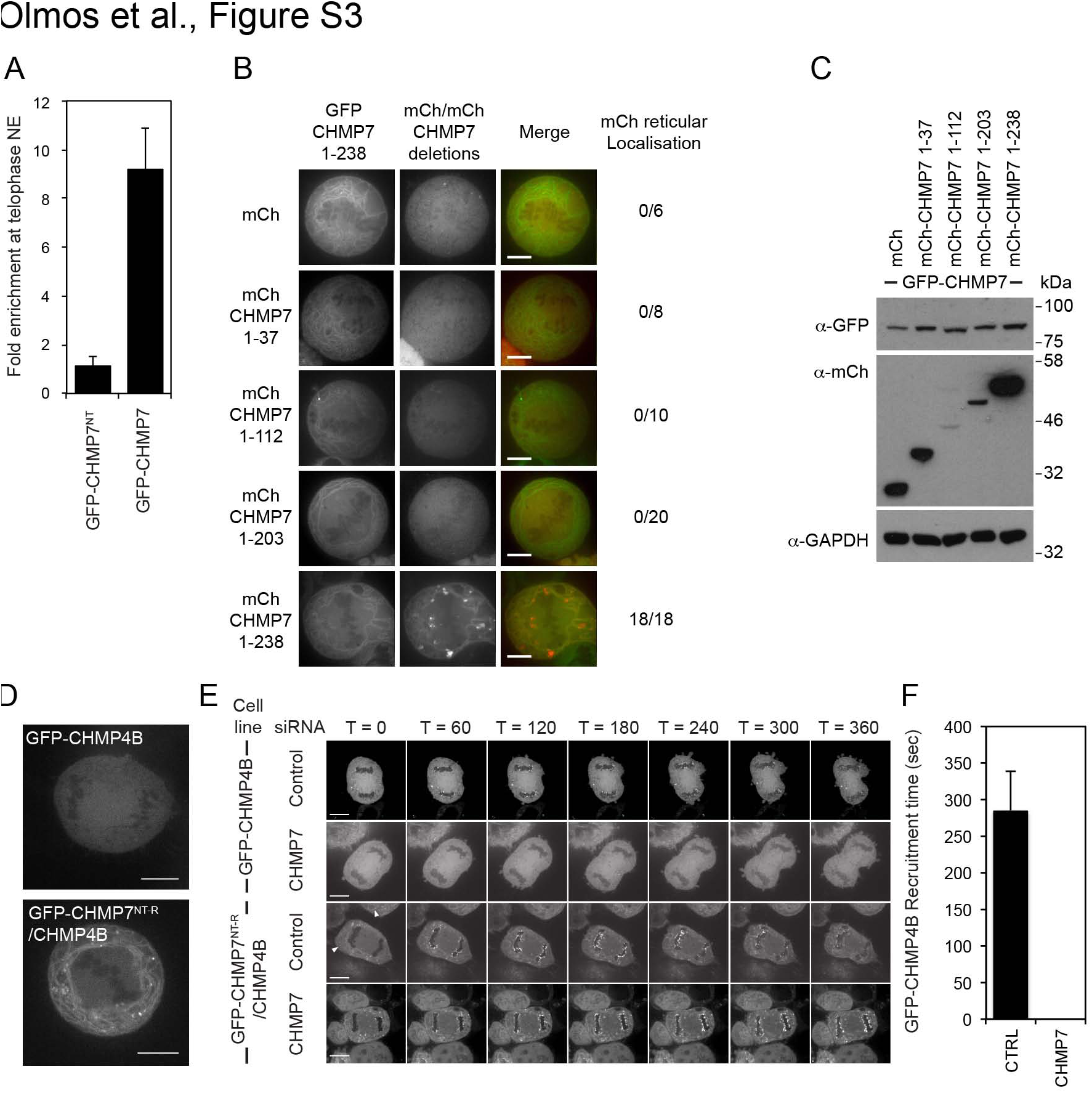
CHMP7^NT^ directs ER localisation and is necessary for subsequent enrichment of CHMPs at the reforming NE. A. HeLa cells stably expressing GP-CHMP7 or GP-CHMP7^NT^ were imaged live and the degree of enrichment at the reforming NE was assessed by quantification of the maximal fluorescence intensity achieved at the NE compared to an adjacent region of ER. B. HeLa cells stably expressing both GFP-CHMP7^NT^ (1-238) and the indicated mCh-CHMP7 deletions were imaged live and reticular localisation in mitotic cells was scored. C. Resolved cell lysates from cells from B were examined by western blotting with anti-mCh, anti-GAPDH or anti-GFP antisera. D. HeLa cells stably expressing GFP-CHMP4B or GFP-CHMP7^NT^/CHMP4B were imaged live during mitosis. Images representative of all cells imaged and 5/5 captured cells per condition. E. HeLa cells stably expressing GFP-CHMP4B or GFP-CHMP7^NT-R^/CHMP4B were transfected with the indicated siRNA and imaged live. Images representative of 3/3 movies (GFP-CHMP4B, Control siRNA), 3/3 movies (GFP-CHMP4B, CHMP7 siRNA), 5/5 movies (GFP-CHMP7^NT-R^/CHMP4B, Control siRNA), 6/6 movies (GFP-CHMP7^NT-R^/CHMP4B, CHMP7 siRNA), arrowheads depict ER localisation. F. Quantification of duration of GFP-CHMP4B localisation from movies in E (Duration presented in seconds ± S.D. Control, n = 6; CHMP7 siRNA n = 7).

**Figure S4:**
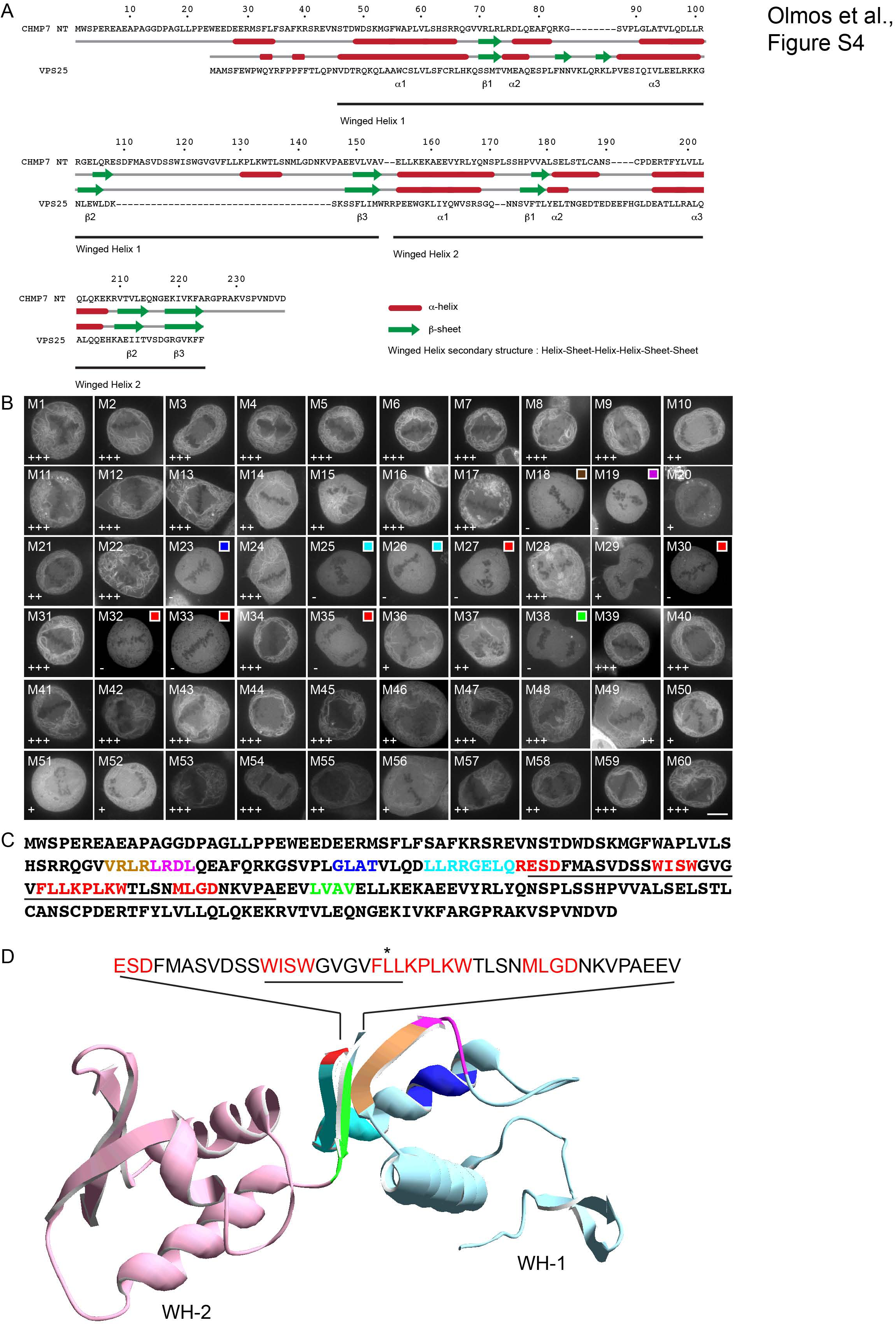
Sequence analysis and mapping of ER-localisation determinants of CHMP7^NT^. A. Manual alignment of CHMP7^NT^ and VPS25. Secondary structural prediction of elements within CHMP7^NT^ using JPred(Drozdetskiy *et al*, 2015) and aligned against the secondary-structural elements obtained from the crystal structure of VPS25 (PDB 3CUQ chain C)(Im & Hurley, 2008). Residue numbering given for CHMP7. B. HeLa cells expressing the indicated GFP-CHMP7^NT^ plasmids were imaged live. Mitotic cells were captured to maximise ER visibility. Scale bar is 10 μm. Images representative of all cells imaged and 3/3 captured images per mutation. Residues mutated in blocks of 4 sequential amino acids were are: M1, W^2^S^3^P^4^E^5^ – AAAA; M2, REAE – AAVA; M3, APAG – VAVA; M4, GDPA – AAAV; M5, GLLP – AAAA; M6, PEWE – AAAA; M7, EDEE – AAAA; M8, RMSF – AAAA; M9, LFSA – AAAV; M10, FKRS – AAAA; M11, REVN – AAAA; M12, STDW – AAAA; M13, DSKM – AAAA; M14, GFWA – AAAV; M15, PLVL – AAAA; M16, SHSR – AAAA; M17, RQGV – AAAA; M18, VRLR – AAAA; M19, LRDL – AAAA; M20, QEAF – AAVA; M21, QRKG – AAAA; M22, SVPL – AAAA; M23, GLAT – AAVA; M24, VLQD – AAAA; M25, LLRR–AAAA, M26, GELQ – AAAA; M27, RESD – AAAA; M28, FMAS – AAAA; M29, VDSS – AAAA; M30, WISW – AAAA; M31, GVGV – AAAA; M32, FLLK – AAAA; M33, PLKW – AAAA; M34, TLSN – AAAA; M35, MLGD – AAAA; M36, NKVP – AAAA; M37, AEEV – VAAA; M38, LVAV – AAVA; M39, ELLK – AAAA; M40, EKAE – AAAA; M41, EVYR – AAAA; M42, LYQN – AAAA; M43, SPLS – AAAA; M44, SHPV – AAAA; M45, VALS – AAAA; M46, ELST – AAAA; M47, LCAN – AAAA; M48, SCPD – AAAA; M49, ERTF – AAAA; M50, YLVL – AAAA; M51, LQLQ – AAAA; M52, KEKR – AAAA; M53, VTVL – AAAA; M54, EQ – AA; M55, NG – AA; M56, EKIV – AAAA; M57, KFAR – AAAA; M58, GPRA – AAAA; M59, KVSP – AAAA; M60, VNDV – AAAA. Images were scored for strength of ER localisation. Colour-coded key to positions of residues on the homology model given for mutations that abolished ER-localisation. C. Sequence of CHMP7^NT^ with mutants (M18, M19, M23, M25, M26, M27, M30, M32, M33, M35, M38) that disrupt localisation to the ER highlighted in colours – mutations in the WH1 β2 - β3 insertion (underlined) are highlighted in red. D. Homology model of CHMP7^NT^ tandem WH-core (amino acids 19-224; δ107-148) based upon VPS25 3CUQ. Modelled structure in blue (WH1) and pink (WH2), position of mutants that disrupt localisation to the ER highlighted in colours from C. Position of insertion between β2 and β3 given as text sequence with δ118-128 region underlined and L127 highlighted. Residues highlighted in red correspond to amino acids that abolished ER localisation when mutated, as depicted in Figure S4B and S4C.

**Figure S5:**
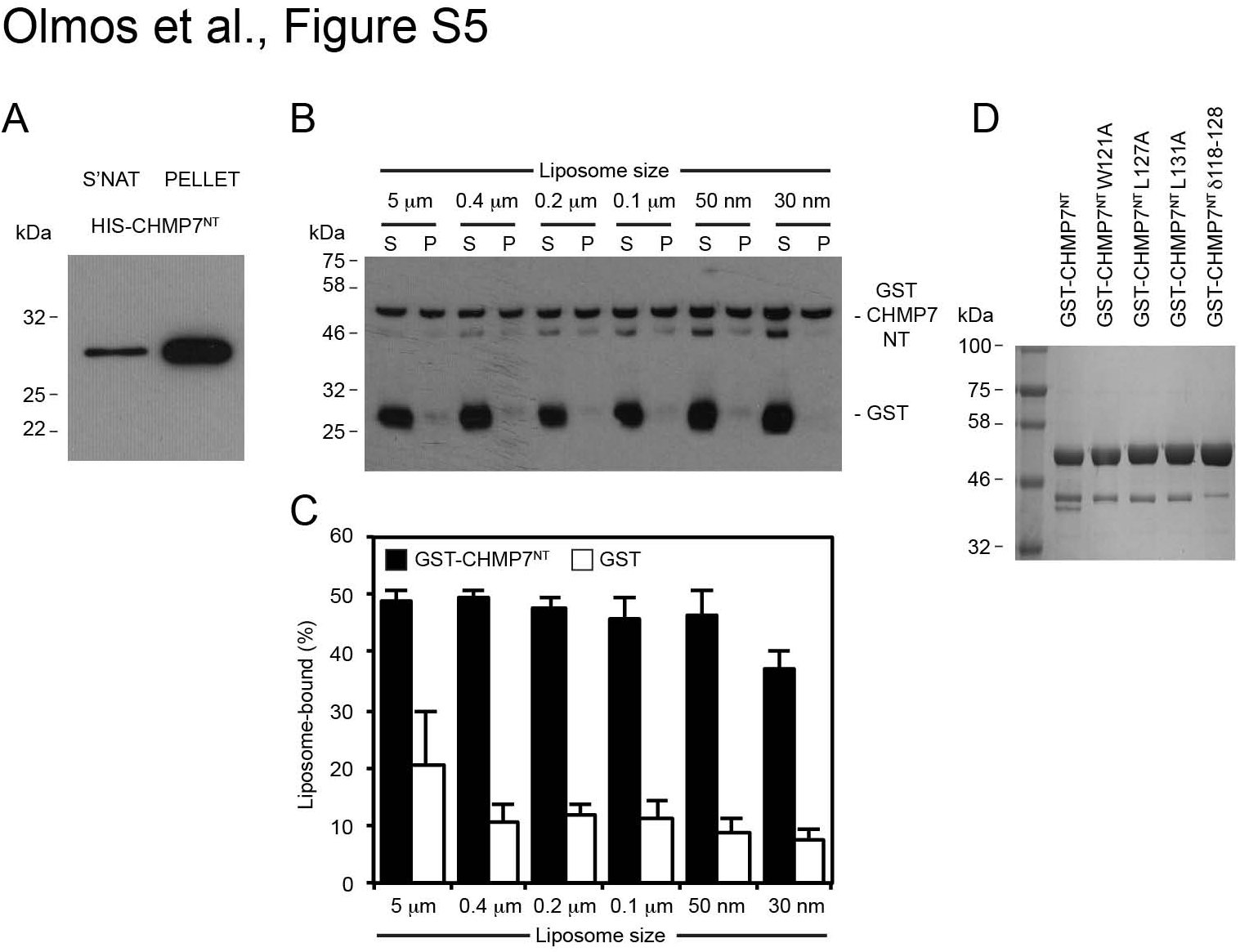
Characterisation of CHMP7 lipid binding. A. Resolved pellet and supernatant fractions of HIS-CHMP7^NT^ interaction with synthetic liposomes (60% DOPC, 20% DOPS, 20% DOPE) were analysed by western blotting with anti-HIS antisera. Blot representative of 2 independent experiments, performed in duplicate and triplicate respectively. B. Resolved pellet and supernatant fractions of GST-CHMP7^NT^ incubated with synthetic liposomes of the indicated size. C. Quantification of binding from Figure S5B. Values quotes as mean ± S.E.M. from 3 independent experiments; two experiments were performed in duplicate and averaged. Statistical significance calculated using 1-way ANOVA with Tukey’s multiple comparison test from experimental means. Significance was not achieved in any case. D. Coomassie stained gel of GST-CHMP7^NT^ mutations from Fig 3E.

## Supplemental Movie Legends

### Movie 1: GFP-CHMP7 localises to the ER and reforming NE

HeLa cells stably expressing GFP-CHMP7 were imaged live, frames were acquired every 30 seconds and displayed at 10 frames per second. Movie representative of 21/21 acquired movies.

### Movie 2: GFP-CHMP7 localises to the ER and reforming NE

Cos7 cells stably expressing GFP-CHMP7 at endogenous levels were imaged live, frames were acquired every 30 seconds and displayed at 10 frames per second. Movie representative of 5/5 acquired movies.

### Movie 3: GFP-CHMP7 localises to the interphase ER

Cos7 cells stably expressing GFP-CHMP7 at endogenous levels were imaged live, frames were acquired every 30 seconds and displayed at 10 frames per second. Movie representative of 72/72 captured live cells.

### Movie 4: GFP-CHMP7 colocalises with ER-markers

HeLa cells stably expressing GFP-CHMP7 were transfected with a plasmid encoding BIP-mCh-KDEL and imaged live, frames were acquired every 30 seconds and displayed at 10 frames per second. Movie representative of 3/3 acquired movies.

### Movie 5: GFP-CHMP7^R^ localises to the ER and reforming NE

HeLa cells stably expressing siRNA-resistant GFP-CHMP7^R^ at endogenous levels were transfected with CHMP7-targetting siRNA and imaged live, frames were acquired every 30 seconds and displayed at 10 frames per second. Movie representative of 3/3 acquired movies.

### Movie 6: GFP-CHMP7^NT^ localises to the ER

HeLa cells stably expressing GFP-CHMP7^NT^ were imaged live, frames were acquired every 30 seconds and displayed at 10 frames per second. Movie representative of 22/22 acquired movies.

### Movie 7: GFP-CHMP7^NT^ colocalises with the ER-markers

HeLa cells stably expressing GFP-CHMP7^NT^ were transfected with a plasmid encoding BIP-mCh-KDEL and imaged live, frames were acquired every 30 seconds and displayed at 10 frames per second. Movie representative of 3/3 acquired movies.

### Movie 8: GFP-CHMP7δNT remains cytosolic during mitosis

HeLa cells stably expressing GFP-CHMP7 δNT were imaged live, frames were acquired every 30 seconds and displayed at 10 frames per second. Movie representative of 5/5 acquired movies.

### Movie 9: GFP-CHMP7 δ118-128 remains cytosolic during mitosis

HeLa cells stably expressing GFP-CHMP7 δ118-128 were imaged live, frames were acquired every 30 seconds and displayed at 10 frames per second. Movie representative of 3/3 acquired movies.

### Movie 10: GFP-CHMP7 W118A remains cytosolic during mitosis

HeLa cells stably expressing GFP-CHMP7 W118A were imaged live, frames were acquired every 30 seconds and displayed at 10 frames per second. Movie representative of 3/3 acquired movies.

### Movie 11: GFP-CHMP7 W121A remains cytosolic during mitosis

HeLa cells stably expressing GFP-CHMP7 W121A were imaged live, frames were acquired every 30 seconds and displayed at 10 frames per second. Movie representative of 3/3 acquired movies.

### Movie 12: GFP-CHMP7 F126A remains cytosolic during mitosis

HeLa cells stably expressing GFP-CHMP7 F126A were imaged live, frames were acquired every 30 seconds and displayed at 10 frames per second. Movie representative of 3/3 acquired movies.

### Movie 13: GFP-CHMP7 L127A remains cytosolic during mitosis

HeLa cells stably expressing GFP-CHMP7 L127A were imaged live, frames were acquired every 30 seconds and displayed at 10 frames per second. Movie representative of 3/3 acquired movies.

### Movie 14: GFP-CHMP7 L131A remains cytosolic during mitosis

HeLa cells stably expressing GFP-CHMP7 L131A were imaged live, frames were acquired every 30 seconds and displayed at 10 frames per second. Movie representative of 3/3 acquired movies.

## Methods

### Cell Culture

HeLa (ATCC), GP2-293 (Clontech) or Cos7 (ATCC) cells were cultured in DMEM containing 10% FBS, Penicillin (100U/ml) and Streptomycin (0.1 mg/ml). Stable cells lines were generated by transduction using MLV-based retroviruses as described previously(Carlton *et al*, 2008), and selected using Puromycin (200 ng/ml), G418 (500 μg/ml) or hygromycin (200 mg/ml) as necessary. Where necessary, cells were sorted to monoclonality by limiting dilution or FACS. Cell lines stably expressing Histone 2B-mCherry (H2B-mCh) and GFP-NLS have been described previously(Olmos *et al*, 2015).

### Plasmids

The human CHMP7 coding sequence was amplified from Image Clone 5551762 (GE Healthcare) and subcloned using in-frame 5’ EcoRI and 3’ NotI restriction sites into the retroviral packaging vectors pCMS28-GFP-EcoRI-NotI-XhoI (ENX; IRES-Puro), pNG72-mCh-ENX (IRES-Neo) or pNG72-ENX (IRES-Neo) (MCS-modified versions of kind gifts from Dr Chad Swanson and Prof Mike Malim, KCL) (Gallois-Montbrun *et al*, 2007). Deletions and mutations of, or addition of epitope tags to, CHMP7 were created by standard PCR-based molecular biology procedures and inserted into pCMS28-ENX, pCMS28-GFP-ENX, or pNG72-GFP-ENX. All constructs were verified by sequencing. CHMP7 was rendered resistant to siRNA oligo-1 through the introduction of silent mutations G217G, E218E, K219K, I220I, V221V, K222K. Coding sequences were cloned EcoRI-NotI into pGEX or pET28a vectors for expression of recombinant proteins.

For retroviral transduction, above constructs in retroviral packaging vectors were transfected with pVSVG into GP2-293 cells (Clontech). Supernatants were harvested, clarified by centrifugation (200 × g, 5 minutes), filtered (0.45 mm) and used to infect target cells in the presence of 8 μg/ml polybrene (Millipore) at MOI < 1. Antibiotic selection was applied after 48 hours.

### Antibodies

An antibody against GAPDH (MAB374) was from Millipore, Calnexin (ab22595) was from Abcam, Tubulin (DM1A) was from Sigma, CHMP2A (104771-AP) was from Proteintech (note, later batches of this polyclonal displayed a non-specific band, marked with an asterisk in the relevant blots), GFP (7.1/13.1) was from Roche, HA.11 was from Covance. Anti-RanBP3 (134052) was from Abcam. Anti-ERp57 (TO2) was from Sigma, anti-EEA1 (C45B10) was from Cell Signaling Technology. Anti-GST (27457701V) was from GE Healthcare. Anti-HIS (2365) was from Cell Signaling Technology. Alexa conjugated secondary antibodies were from Invitrogen and HRP-conjugated secondary antibodies were from Millipore.

### Subcellular Fractionation

Following the method of Graham, 2002 (Graham, 2002), cells (approx. 100 million) were collected and swollen for 10 minutes in homogenisation buffer (0.25 M Sucrose, 1 mM EDTA, 10 mM Hepes (pH 7.4)) and broken by 10 passages through a 12 μm-spaced ball-bearing homogeniser (Isobiotec). Nuclei and cellular debris were pelleted by centrifugation (10 minutes at 1700 × g) and a post-nuclear supernatant was layered on top of a 13ml continuous (0-25%) iodixanol gradient atop a 50% iodixanol cushion. The gradient was centrifuged at 150,000 × g for 15 hours using an SW28 Ti rotor (Beckmann) and 0.5 ml fractions were collected for analysis by SDS-PAGE and western blotting.

### SDS-PAGE and western blotting

Cell lysates and fractions were denatured in Laemmli buffer and resolved using SDS-PAGE. Resolved proteins were transferred onto nitrocellulose by western blotting and were probed with the indicated antisera in 5% milk. HRP-conjugated secondary antibodies were incubated with ECL Prime enhanced chemiluminescent substrate (GE Healthcare) and visualized by exposure to autoradiography film.

### Transient transfection of cDNA

HeLa cells were transfected using Lipofectamine-3000 (Life Technologies) according to the manufacturers instructions. 293GP2 cells were transfected using linear 25-kDa polyethylenimine (PEI, Polysciences, Inc.)

### siRNA transfections

HeLa cells were seeded at a density of 1E5 cells/ml and were transfected with siRNA at 20 nM, 2 hours after plating using RNAi-MAX (Invitrogen), for 72 hours. The following targeting sequences that have already been demonstrated to achieve potent and specific suppression of the targeted CHMP were employed: Control – Dharmacon Non-targeting control D-001810, CHMP7-1 (GGGAGAAGATTGTGAAGTTdTdT(Morita *et al*, 2011), CHMP7-2 (GGAGGUGUAUCGUCUGUAUdTdT, M-015514-11). Given the similarity between the CHMP7 sequence targeted by oligo-1 and RanBP3, we ensured that the CHMP7 oligos used in this study did not suppress endogenous RanBP3, whereas a RanBP3-targeting siRNA (Smartpool M-011484) effectively suppressed endogenous RanBP3 (Figure 1E).

### Production of recombinant proteins

BL21 (DE3) * *E. coli*, expressing plasmids encoding GST-or HIS-tagged proteins, were resuspended in bacterial lysis buffer (20mM Hepes (pH 7.4), 500 mM NaCl, 3.5% glycerol and supplemented with Complete mini, EDTA-free protease inhibitor (Roche) and 1 mM PMSF. Cells were lysed by addition of lysosyme (1 mg/ml, 15 minutes), Triton X100 (0.25%, 15 minutes) and were snap frozen in liquid nitrogen. Cells were thawed on ice, clarified through addition of DNAse1 (20 μg/ml) and soluble proteins were collected by centrifugation at 28000 × g for 30 minutes. Proteins were immobilised on Glutathione Sepharose 4b or Ni-NTA agarose resins, washed extensively in wash buffer (20mM Hepes, pH 7.4, 150mM NaCl, 3.5% Glycerol). Proteins were eluted from Glutathione Sepharose 4β resin in wash-buffer supplemented with 10mM reduced Glutathione (pH 8) and were dialysed against wash-buffer. Eluted proteins were stored at −80 °C. HIS-tagged proteins expressed from pET28a were expressed and harvested similarly, barring that all buffers contained 20 mM imidazole, and that proteins were eluted with a step gradient of imidazole.

### Liposome binding assays

Liposome binding assays were performed as previously described(Cozier *et al*, 2002). Briefly, Folch extract was resuspended at 10mg/ml in CHCl_3_:MeOH (19:1) and was dried as a film onto a round-bottomed glass tube by overnight rotary evaporation. The lipid film was rehydrated at 10 mg/ml under rotation in sucrose buffer (200 mM Sucrose, 20 mM KCL, 20 mM Hepes, pH 7.4) for 1 hour. Alternatively, synthetic liposomes were prepared by drying mixtures of 1,2-dioleoyl-sn-glycero-3-phosphocholine (PtdCho (60%)), 1,2-dioleoyl-sn-glycero-3-phosphoserine (PtdSer (20%)) and 1,2-dioleoyl-sn-glycero-3-phosphoethanolamine (PtdEto (20%)) and resuspending similarly. 2.5% 1-2-dioleoyl-sn-glycerol (DAG) or 1,2-dioleoyl-sn-glycero-3-phosphate (PtdOH) was added if required. Synthetic lipids were from Avanti Polar Lipids. Insoluble matter was removed by centrifugation (1000 × g, 1 minute) and liposomes were generated by bath sonication (5 minutes). In Figure S5B and S5C, liposomes were generated by extrusion of the rehydrated synthetic lipids though indicated defined pore-size nitrocellulose filters (Whatmann) using an Avanti Mini-Extruder. Proteins were diluted to 7 μg/ml in osmotically-matched protein dilution buffer (20 mM Hepes, 120 mM NaCl, 1 mM EGTA, 0.2 mM CaCl_2_, 1.5 mM MgCl_2_, 1 mM DTT, 5mM KCl, pH 7.4, 1% BSA was added to enhance solubility) and were pre-cleared by ultracentrifugation at 120,000 × g for 45 minutes using a TLA100.3 rotor. 1 ml of protein mixture was then combined with 10 μl of liposomes and incubated with shaking at 30 ^o^C for 5 (Figure 3A) or 15 (Figure 3B) minutes. Liposomes were recovered by ultracentrifugation (120,000 × g for 30 minutes); supernatant and pellet fractions were resuspended in equal volumes of Laemmli buffer and analysed by western blotting. Band intensities were quantified by densitometry using ImageJ and liposome-bound fractions were calculated.

### Fixed cell imaging

Cells were imaged using Nikon Eclipse microscopes teamed with confocal (CSU-X1 Andor Spinning Disc/Ixon3 EM-CCD) imaging systems. Images were processed in NIS Elements and exported to Photoshop for assembly into figures. HeLa cells were fixed in MeOH (for CHMP2A-staining) or 4% PFA and subject to processing for immunofluorescence as described previously (Carlton *et al*, 2012).

### Live cell imaging

HeLa cells stably expressing the indicated proteins were plated in 4- or 8-chamber Stickyslides (Ibidi) adhered to a glass number 1 coverslip and transfected with the indicated siRNA where necessary. For analysis of GFP-CHMP7 recruitment, cells were transferred to a inverted spinning disc confocal microscope (Nikon Eclipse, teamed with CSU-X1 Andor Spinning Disc with Ixon3 EM-CCD) with attached environmental chamber and imaged live using a 100x oil-immersion objective, acquiring frames every 30 seconds. For enrichment of GFP-CHMP7 on the forming NE, background-corrected maximal fluorescence fluorescence intensities on the telophase NE were normalised against those on regions of adjacent ER.

For analysis of nucleo-cytoplasmic compartmentalisation, as described in(Olmos *et al*, 2015), cells were synchronised using a double thymidine block and 54 hours after siRNA transfection (10.5 hours after release from the second thymidine block), cells were transferred to a inverted spinning disc confocal microscope with attached environmental chamber and imaged live for 4 hours using a 20x dry objective and a 1.5 × magnification lens, acquiring frames every 1-5 mins. The ratio of background-corrected, area-normalised, GFP-positive pixel intensities within the cytoplasm and mCh-H2B demarcated nuclei at the indicated intervals were obtained using NIS-elements. Typically 20 daughter cells per siRNA treatment were analysed and the indicated number of independent experiments were performed as described in the relevant figure legends.

### Modelling

CHMP7 residues 1-238 comprising tandem WH domains identified by HH-Pred (with a 4 residue flexible linker replacing the insertion (residues 107-148)), was submitted to Swiss-Model server using a template-directed homology search, returning VPS25 (3CUQ). Models were built and exported from Swiss PDB-viewer.

### Statistical analysis

2-tailed Student’s T-tests, or ordinary 1-way ANOVA with the indicated post-hoc tests were used to assess significance between test samples and controls and were performed using GraphPad Prism. N-numbers given as the number of independent experiments, n-numbers given as the number of cells analysed.

